# Specialists drive biodiversity scaling in symbiotic relationships

**DOI:** 10.64898/2026.03.13.711632

**Authors:** Colin J. Carlson, Jeremy B. Yoder, Timothée Poisot

## Abstract

The stunning diversity of symbiotic life forms, and their unique vulnerability to extinction, emerge from the close relationship between host and symbiont species richness. The general form of this relationship should be linear, but simulation studies have shown that it becomes sub-linear (and even power law-like) when sampling within a host-symbiont network. Here, we resolve this paradox with a new mathematical model of scaling in bipartite graphs, based on the independent behavior of specialist (single-host) and generalist (multi-host) symbionts. Using this model, we show that specialists constrain the architecture of ecological networks, and at global scales, drive the accumulation of symbiont biodiversity. By definition, specialists also face the highest risk of coextinction with their hosts and — despite substantial uncertainty about their true richness — we show that they could easily account for the majority of threatened species on Earth. Our study reveals that symbiosis remains one of the most poorly-understood building blocks of ecosystem function and evolutionary diversification, and serves as a reminder that foundational macroecological principles are still waiting to be discovered from first principles.

---

> *We are astonishingly ignorant about how many species are alive on earth today, and even more ignorant about how many we can lose yet still maintain ecosystem services that humanity ultimately depends upon*. — Robert May, 2011 *(1)*
>
> *Small amounts, adding up to make a work of art*. — Stephen Sondheim, 1984 (*2*)

Symbiosis is a close ecological relationship between species that is typically beneficial for one (parasitism and commensalism) or both (mutualism) (*3*). These relationships are ubiquitous across ecosystems and evolutionary history, and can create dependencies between profoundly different organisms, such as acacia trees and the ants that guard them from herbivores, or spotted salamanders and the green algae that colonize their cells (*4*). In many cases, one species (the host) provides space or food that another (the symbiont, also sometimes called a dependent or affiliate species) cannot survive without. Human-caused environmental change can disrupt these fragile interactions, and even lead to the coextinction of hosts and their symbiont community (*5, 6*); the downstream consequences for ecosystem function and stability are poorly understood, but generally agreed to be far-reaching and potentially dire (*7–10*).

The closeness of these ecological relationships suggests that there might be a similarly close relationship between symbiont and host biodiversity. Over time, symbionts have diversified alongside their hosts through both co-divergence and host switching (*11–13*); over space, symbiont and host species richness are often closely correlated (*14–18*); and, in the face of biodiversity loss, higher host extinction rates are expected to result in higher symbiont coextinction rates (*5, 6*). Despite all this, there is no canonical ecological law that describes how host and symbiont diversity scale with each other. Previous studies agree that this relationship should be driven by the host specificity of the symbionts, but disagree about whether the resulting pattern is linear (*19, 20*) or sub-linear (*5, 21, 22*). Here, we derive not one but two general forms of this relationship that resolve this paradox. Using insights from these mathematical models, we show how specialists — symbionts that depend on a single host species — drive the structure of ecological networks, the scaling of symbiont biodiversity, and the vulnerability of symbiotic interactions to biodiversity loss.

## The scaling paradox

By definition, the diversity of host-specific symbionts must scale ***linearly*** with the diversity of their hosts, with a slope given by the mean per-host symbiont richness. In the 1980s, this property was applied to beetles and the trees they feed upon to generate some of the first estimates of global arthropod diversity (*23–25*). Later work showed that this relationship can be generalized to all symbionts simply by dividing by the mean symbiont host range. For example, Poulin and Morand collected estimates of these two parameters from the literature and used them to produce the first estimate of the global diversity of parasitic worms of vertebrates (*19*).

In 2014, Strona and Fattorini revisited these estimates using simulation methods, and discovered that *within* a given sample, parasitic worms appear to scale ***sub-linearly*** with hosts (*21*). Just as species discovery rates decrease over larger geographic areas (*26, 27*), each additional host sampled is likely to have fewer unique parasites. Strona and Fattorini hypothesized that this scaling relationship — like the species-area relationship, and so many others in biology (*26, 28–30*) — might be adequately described by a power law (*S* ∼ *H* ^*z*^, 0 ≤ *z* ≤ 1). More recently, we found that this pattern is widespread across different types of symbiosis, and used power scaling laws to re-estimate the global diversity of mammalian viruses and vertebrate parasites (*22, 31*). The idea that this relationship takes the form of a power law was appealing, given the ubiquity of power laws in the biological sciences, but we were unable to provide any mechanistic basis for this hypothesis.

### The symbiont species-host relationship

Here, we describe two alternate solutions — one so-far overlooked, and one entirely novel — that better explain the scaling of host and symbiont biodiversity from first principles, and reconcile the linear and sub-linear aspects of this relationship.

#### Symbiont specialization drives subgraph scaling

The first analytic approximation of symbionts’ scaling with host diversity was proposed by Koh, Colwell, and colleagues more than 20 years ago, when they repurposed an unbiased estimator that was developed for rarefaction curves (i.e., species accumulation over a sampling unit, such as quadrats) (*32*), and applied it to coextinction rates (*5*). This approach starts from a known network of *H* hosts and *S* symbionts, and asks how many symbionts *s*(*h*) will be left in a network of *h* < *H* hosts (i.e., with a host extinction rate of *E*_*h*_ = 1 − *h*⁄*H*). For a symbiont with *k* hosts in the full network, the probability of observing 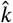 hosts (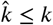) in the sample of *h* hosts is given by:

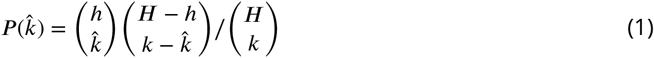

The probability that a given symbiont will have zero remaining hosts in the sample (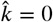), and so will have gone extinct along with its hosts, is given by:

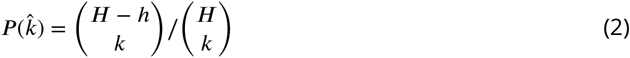

Therefore, if *s*_*k*_ is the number of symbionts with *k* hosts, then the total expected number of coextinctions is given by 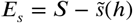, where:

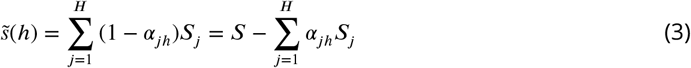

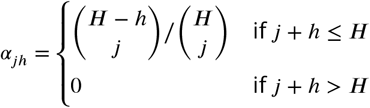

With knowledge of the global diversity of hosts (*H*) and symbionts (*S*), and the distribution of symbiont host range (*S*_*j*_), coextinction rates can be precisely estimated using equation 3, without a need for the kinds of simulations that are commonplace in coextinction studies.

The Koh-Colwell method shows that the scaling of symbiont and host diversity is, essentially, singularly driven by symbiont specialization. However, it also requires perfect knowledge of the full global network (*S* and *S*_*j*_), and can only be used to travel “down the curve” to estimate *s*(*h*) for values of *h* < *H*; equation 3 cannot be used to extrapolate *S* from *h, s*, and *H* (but see “How many symbionts are there?” below). Therefore, this approach correctly reproduces, but does not intuitively explain, the sub-linear scaling of symbiont and host species richness observed in simulated subgraphs. Similarly, it provides an analytical shortcut to estimate coextinction rates without simulation, but requires nearly as much information as those simulations.

#### Symbiont species richness scales linearly *and* sub-linearly

Here, we derive an alternative, closed-form approximation of symbiont scaling that relies on just three network-level parameters, which can be estimated from relatively small samples of the full network. The relationship between host and symbiont species richness (*H* and *S*) always follows an identity that emerges from the number of links (equivalent to the number of edges or associations) in the network, a property first used by Poulin and Morand to estimate global parasite diversity (*19*):

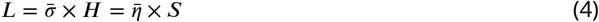

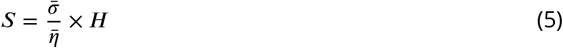

where 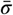 and 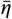 are the average number of symbionts associated with each host and vice versa. This linear scaling is reproduced when subgraphs are sampled by edges: for a given sample of *l* randomly chosen links, the subgraph should include roughly 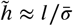 hosts and 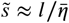 symbionts.

How, then, does sampling by hosts produce a sub-linear relationship? Within the global network, *specialist* symbionts (here defined as those with a single host: *η*_*s*_ = 1; also sometimes referred to as *monoxenous* in the case of parasites) and *generalist* symbionts (two or more hosts: *η*_*g*_ > 1; i.e., *stenoxenous*) form two distinct subgraphs that share some hosts, but no symbionts. These subgraphs scale separately, but must still follow equation 5 (**Figure 1**):

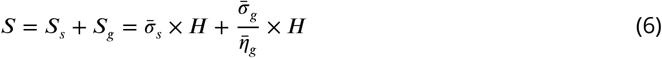

**Figure 1.**
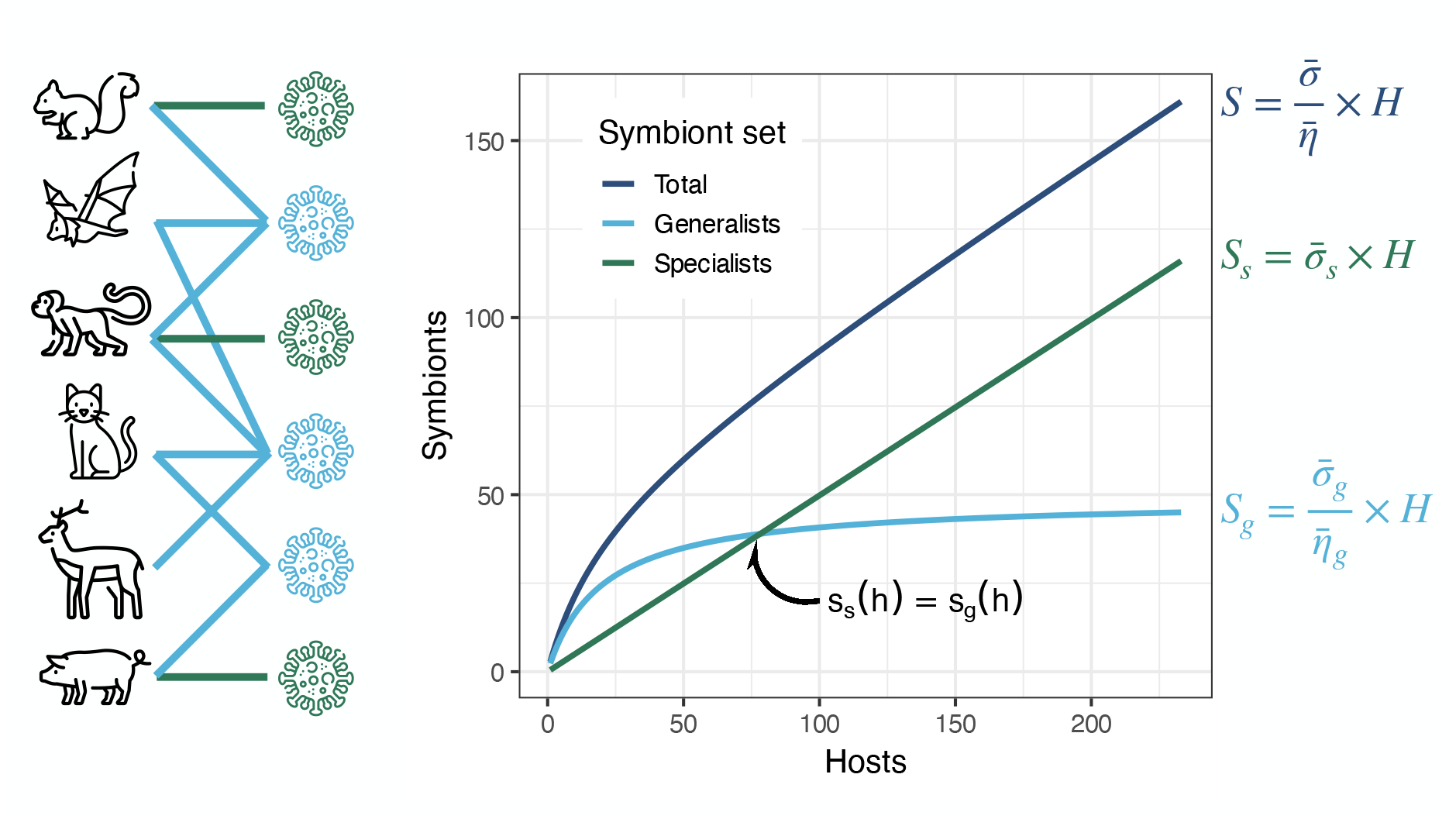
Specialists scale linearly, while generalists scale asymptotically. In the network of associations between hosts and symbionts (mammals and viruses, left), the relationship between host and symbiont diversity is asymptotic for generalists (right, light blue line) but linear for specialists (right, green line); the sum of these relationships scales sublinearly with host diversity (right, dark blue line, equation 6). Past the point at which generalist symbiont diversity is equal to specialist symbiont diversity (callout), the majority of symbionts are specialists.

This property applies to any subgraph of *h* hosts (*h* < *H*) and all of their associated *s* symbionts:

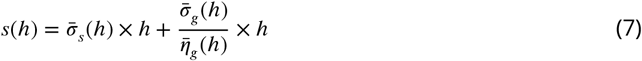

However, *s*(*h*) does not scale linearly with *h*. Every host added to the network has the same expected number of specialists (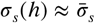) and generalists (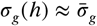). However, as hosts are added to the network, the observed host range of generalists expands roughly linearly (**Figure S1**):

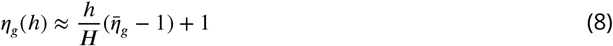

Therefore, the overall curve can instead be approximated by:

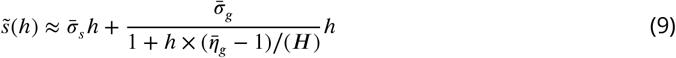

The result is intuitive: every host added to the network may bring a new set of specialists, but the average number of new generalists added with each host gradually approaches zero. Superim-posed, this linear scaling of symbionts and asymptotic scaling of generalists produces a sub-linear curve without an easily-identified pattern (**Figure 1**).

Previously, we showed that this curve could be approximated by a power law fit (*s*(*h*) ∼ *h*^*z*^; 0 ≤ *z* ≤ 1), where a fully-connected network would necessarily scale flatly (*z* = 0), and a fully-specialist network would scale linearly (*z* = 1); most real networks fall somewhere in-between, with the exponent *z* constituting a sort of global measure of host specificity. To compare the power law method and closed-form solution, we applied both to a set of 253 real bipartite ecological networks (primarily plant-pollinator, plant-seed disperser, and host-parasite networks; see **Methods**). In a simple linear model, the proportion of specialist symbionts (*S*_*s*_⁄*S*) and the average host range of generalists (*η*_*g*_) are able to explain a combined 91% of variation in estimated values of *z* (**Figure S2**). For any given network, the three-parameter approximation can produce an equally good fit to the power law, without the assistance of simulation *or* curve-fitting methods — though the Koh-Colwell estimator (which uses information on the full network) consistently outperforms both (**Figure S3**).

### Do specialists structure ecological networks?

If specialists scale “separately” from generalists, could they also have distinct effects on network architecture? By partitioning symbiont communities into specialists and generalists, we effectively create two distinct subgraphs of symbiont-host interactions, which we hypothesized might have unique structural constraints. For example, the connectance of a network — the proportion of realized links out of total possible links — is given by:

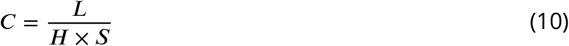

In the specialist-host network, each specialist has exactly one link (*L*_*s*_ = *S*), while each generalist must have at least two or more links (*L*_*g*_ ≥ 2*S*_*g*_). These set the minimum connectance of the total network, which cannot be less than *C*_*s*_ = 1⁄*H*. While generalists can be fully connected (*L*_*g*_ ≤ *S*_*g*_ × *H*_*g*_), specialists also introduce *F* = *S*_*s*_(*H* − 1) forbidden links, creating a maximum possible connectance. Therefore, specialists set hard constraints on network connectance:

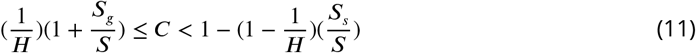

Specialists could just as easily have a similar effect on other commonly used metrics of ecological network structure, such as modularity or nestedness. If so, widely-known patterns in network ecology might be, to borrow very loosely from Gould and Lewontin (*33*), “spandrels”—that is, neutral patterns created by the structural constraints that are introduced by the mere presence of specialists in the network, rather than true reflections of the higher-order structure of the full community.

To test this hypothesis, we compared three measures of network architecture across the same real ecological networks (see **Methods**). Unsurprisingly, we find that networks with a higher proportion of specialists (*S*_*s*_⁄*S*) are significantly less connected, more modular, and less nested (**Figure 2A**). If specialists are removed, these networks become equally or more connected (by definition), and with very rare exceptions, less modular and more nested; network metrics for the full and generalist-only networks are generally correlated, but much more weakly than expected (**Figure 2B**). Moreover, the proportion of specialists is still correlated with the connectance, modularity, and nestedness of the generalist-only networks, but substantially less so, without conspicuous corners of implausible parameter space (see **Figure S4**). Taken together, these findings suggest that these relationships are not merely correlations between network metrics and the full symbiont degree distribution, but rather, true structural constraints created specifically by specialists.

**Figure 2.**
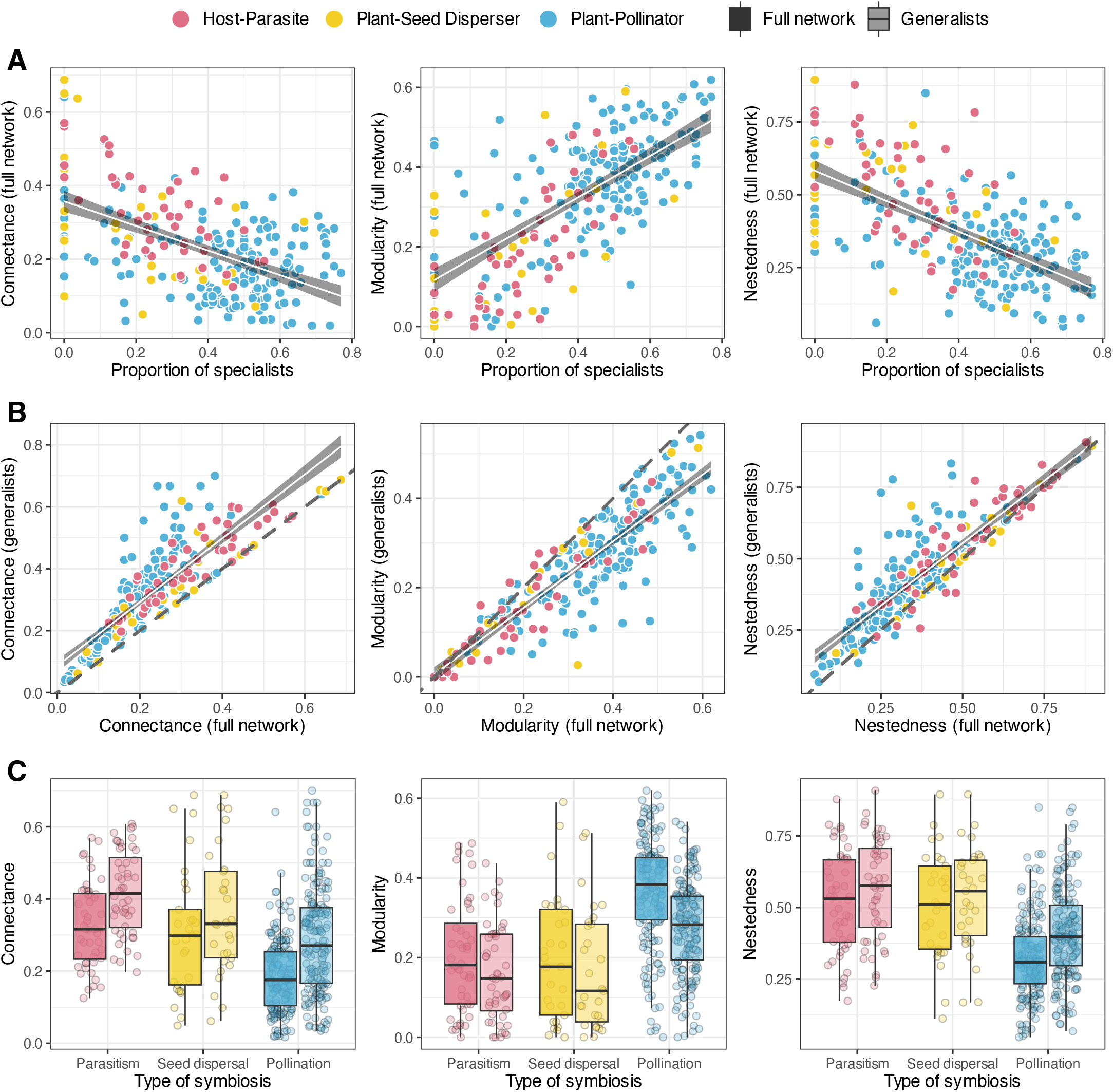
Specialists create spandrels in network architecture. Each point represents one of 246 species interaction networks (see **Methods**). (A) The proportion of specialists constrains commonly-used metrics of network architecture. After removing specialists, these metrics can change substantially (B), and differences between ecological interaction type persist, but become less pronounced (C).

Do these constraints confound ecological inference? For example, plant-pollinator networks are less connected, more modular, and less nested than other network types; these patterns could be driven by a large number of singleton interactions, which are common in short-term, site-specific datasets of floral visitors. To test this hypothesis, we tested whether differences in these metrics persist after removing specialists **(Figure 2C, Table S1**). For all three metrics, we find significant group differences among symbiosis types, and significant effects of removing specialists, but no significant interaction between those factors (i.e., inferred differences in network structure persist even after accounting for the impact of specialists). Based on this finding, we suggest that commonly-used metrics are highly sensitive to the real and structural impact of specialists, but as one among many axes of ecologically-meaningful variation in network architecture.

### How many symbionts are there?

Some symbiotic taxa are relatively well-inventoried: for example, just shy of 1,000 species of ticks (order Ixodida) are known today, and that number has remained mostly stable over the last few decades (*34, 35*). Others are more diverse, but well-enough sampled to generate traditional asymptotic estimates of species richness. For example, there are roughly 300,000 species of flowering plants pollinated by animals (≈88% of flowering plant species), and at least 350,000 described species of arthropods that visit flowers (≈29% of arthropod species) (*36, 37*).

But the most hyperdiverse clades — among them, viruses (*22, 38–41*), parasitic worms (*20, 21, 31, 42*), and parasitoid wasps (*43–45*) — are so speciose, and so poorly described, that they may well be the most uncertain slice in the “pie of life” (*42, 46*). For these groups, the easiest way to estimate their diversity would be to extrapolate from their better-sampled hosts (*19*). Previously, we used a power law approximation to estimate the global diversity of viruses (*22*) and parasitic worms (*31*). Based on our findings so far, we suggest that there is no longer a justification to continue using this method — but as we show below, it is difficult to identify a workable alternative.

#### Existing approaches are mostly unsuitable

Our newly-identified general form for the symbiont species-host relationship does little to enable global estimates of symbiont biodiversity. For values of *h* >> *H*, the approximation given in equation 9 converges on a linear relationship driven entirely by specialists. This is probably true in some sense — approaching global scales, we should expect most “new” symbionts to be specialists — but is a poor basis for extrapolation, particularly given the problem that, in an incomplete sample, many generalist symbionts may be observed on only a single host, and misclassified as specialists (see “Smaller samples have more false specialists” below).

The Koh-Colwell solution appears more promising, given that it was originally developed for sample-based rarefaction. Koh and colleagues did propose that methods designed for extrapolating species richness from incidence data could be similarly repurposed (*5, 32*), but they did not implement their idea because, at the time, it was computationally prohibitive. Over the last two decades, the underlying logic of equation 3 has been extended substantially (*47, 48*), and these methods are now easily implemented (e.g., with the `iNEXT R` package (*49*)). For very well-sampled host communities, these methods could be appropriate. However, as a rule, these approaches should not be used for *h* >> 2*H* (*48*); over larger scales of extrapolation, work by Haegeman and colleagues has shown that any given sample (i.e., small observed network) could be compatible with true communities that range in size by several orders of magnitude (*50*).

Finally, and most fundamentally, inference based on empirical interaction datasets is limited by their incompleteness. This is a key difference from the simulation method we show in **Figure 1**, where each subgraph includes all symbionts associated with the sampled hosts. In contrast, real-world datasets are characterized by imperfect detection of both symbionts and associations (*51*– *53*). If a sizable proportion of hosts have been sampled, and a small proportion of their symbiont community has been characterized, this will invariably lead to substantial underestimates of symbiont diversity. For example, the most comprehensive available dataset on mammalian viruses captures 5,947 virus taxa associated with 1,507 host species (*54*). But whereas humans (the best-inventoried species) have 840 known viruses, the average mammal species in the dataset only has a mean of 8.7 viruses, and roughly one-third of hosts only have a single known virus. It is technically possible to apply standard rarefaction methods to this dataset (*49*) to produce an extrapolated estimate of 18,038 total viruses (range: 17,462–18,614) associated with the 6,759 mammal species worldwide; but realistically this barely provides an informative lower bound on true diversity.

#### The linear method, reconsidered

In lieu of a way to extrapolate from equation 3 or 9, we return to the linear identity given in equation . 5 If 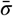 and 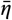 can be reliably estimated, equation 5 can be used to extrapolate *S* linearly from *H*, despite the sublinear scaling of host and symbiont diversity up to that point. The challenge, then, is finding a way to generate reliable estimates of 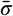 and 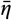: many studies have estimated these parameters from parasite species checklists (*19, 55, 56*), but this approach has an important limitation. In an idealized simulation of “perfect” sampling, where all symbionts are identified when a new host is sampled, estimates of per-host symbiont richness converge on the network-wide average (i.e., 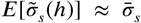 and 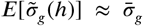), but observed host range increases linearly with sampling (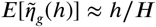) (**Figure S1**). In real-world datasets, hosts are sampled imperfectly: over time, new hosts are sampled, new symbionts are discovered, and new associations are identified between existing hosts and symbionts; as a result, both 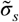 and 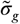 may be unstable through time, without clear convergence on true values of 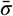 and 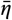.

To illustrate this problem, we compared estimates compiled by Poulin and Morand for different pairs of vertebrate class and parasite type (*19*), to new estimates derived from a global database of animal-helminth associations (*31, 57*), which captures 51,274 interactions between 15,817 parasite species and 6,132 vertebrate host species (∼10% of global vertebrate diversity). Comparing these sources, we find that group differences have mostly been preserved through time, but as more host-specific parasites have been discovered and described (*31*), parasite richness per-host has risen and average host range has decreased sharply (**Figure S5**). Estimates based on more recent data are probably closer to the underlying truth, but they still reflect a small, unrepresentative sample — and are compatible with a wide range of plausible global networks that could vary dramatically in size and structure (similar to (*50*)). As more parasites are described, these values will probably continue to shift. We therefore suggest that linear extrapolation should only be used when parameter estimates can be carefully justified. Next, we discuss one such example.

#### Global viral diversity, revisited (again)

Viruses may well be the most diverse form of life on Earth. Previously, Anthony (*40*), Carroll (*41*), and colleagues developed a two-step method to estimate viral diversity (here meaning operational taxonomic units that are as aligned as possible with International Committee on the Taxonomy of Viruses [ICTV] species definitions). First, they conducted a comprehensive metaviromic inventory of two animal host species; estimated the true number of viruses present in the samples using traditional asymptotic estimators; and finally, estimated per-host viral diversity for the “average” viral family. Second, they extrapolated over 5,291 total mammal species and 25 viral families relevant to human health, and estimated that there are at least ∼1.5 million virus species.

This approach leveraged a unique data source to estimate 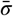 comprehensively, albeit with just two replicates. However, their estimates also made the stunningly unrealistic assumption that all viruses are host-specific (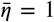). Here, we revisit these estimates, using the most comprehensive source of information on mammal-virus associations (*54*) to calculate 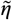 at the viral family level. For several decades, the average host range of known mammal viruses increased sharply, suggesting that our estimates would be highly unstable; but in the last two decades, the emergence of large-scale sequencing-driven wildlife virology — particularly after the 2002 SARS-CoV outbreak (*53*) — has reversed this trend, with a gradual plateau around ∼2.5 known hosts per virus (**Figure S6**).

We calculate average values of 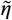 at the family level, making a liberal estimate based on all viruses, and a conservative estimate based only on the host range of ICTV-ratified viruses, which have a more consistent rank and are likely to have their ecology known in greater detail (see **Table S2**). We combine these data with values of 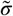 from the original Carroll metagenomic data, but use an updated updated estimate of 6,759 mammal species (*58*), and remove picobirnaviruses, which account for 120 of the estimated 180 viruses associated with the rhesus macaque (*Macaca mulatta*); these viruses have never been confirmed to be vertebrate-infective, and are now suspected to be bacteriophages (*59*). Family-level values of 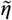 and 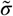 are again averaged to create an “average viral family,” and for consistency, we estimate diversity for 25 priority viral families.

Based on this approach, we estimate there are at least 272,895 to 390,106 mammalian viruses from these 25 viral families (**Figure 3**). As a sensitivity analysis, we separate this estimate for 17 RNA virus families and 8 DNA virus families. Several studies have previously shown that RNA viruses generally have a broader host range than DNA viruses, likely due to their higher mutation rates (*22, 60, 61*). In our previous work using the power law method, separating RNA and DNA viruses led to a substantially higher estimate of total diversity (*22*). As expected, we find that DNA viruses scale much more steeply with host diversity (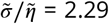 to 3.50) than RNA viruses (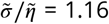 to 1.52). Separating the analysis leads to a estimate of 123,924–189,122 DNA viruses; 133,722–174,197 RNA viruses; and a similar total estimate of 257,646–363,319 mammalian viruses in priority families.

**Figure 3.**
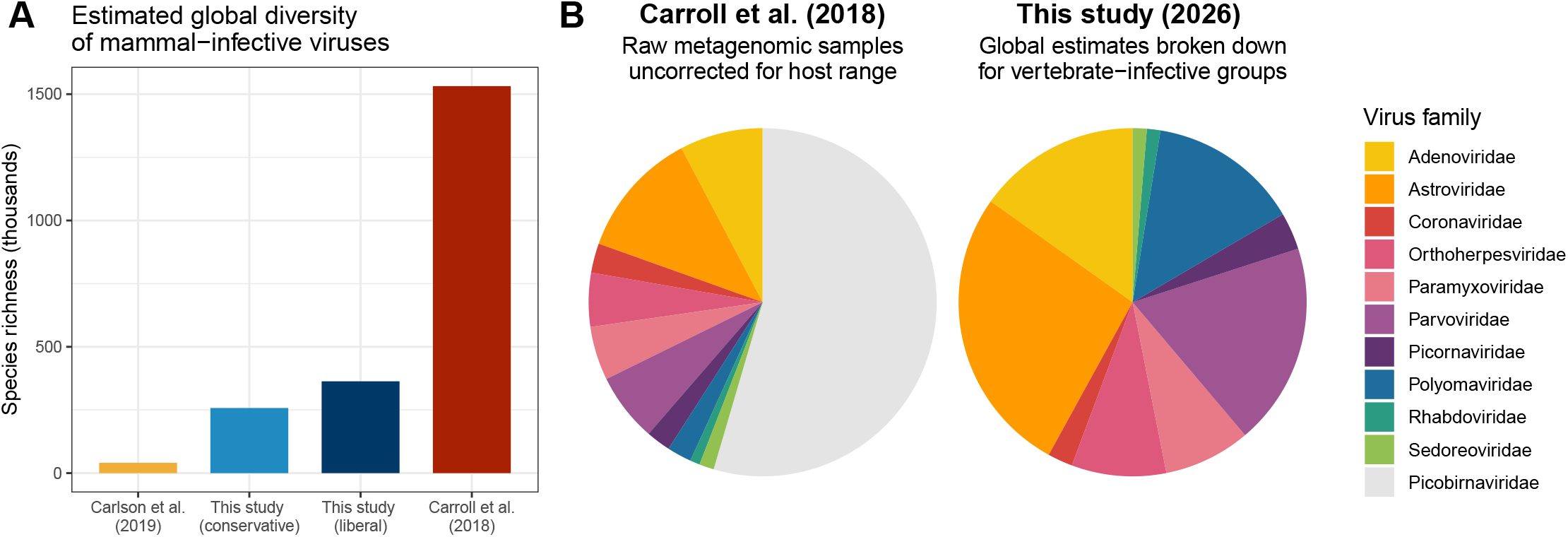
Revised estimates of viral diversity in mammals. (A) This study’s estimates fall between Carroll’s previous estimates, based on linear extrapolation as though all viruses are host-specific (*41*), and our previous estimates, which extrapolated based on an estimated power law scaling (*22*). (Estimates shown are based on the separate calculation for RNA and DNA viruses.) (B) The distribution of viral diversity across 11 viral families in the raw metagenomic data used by Carroll, versus the estimated diversity of these groups based on Table S2.

#### Are most symbionts specialists?

The linear accumulation of specialists can quickly outpace the asymptotic scaling of generalists. Based on equation 9, specialists outnumber generalists globally if:

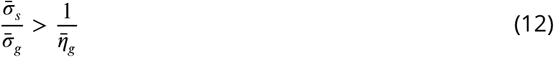

Counterintuitively, this shows that specialists can be globally common even if they appear locally rare: for example, if every generalist has exactly two hosts and no more (i.e., 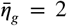, the lowest possible value), then at the level of the average host species, there must only be at least half as many affiliated specialists as generalists. At more realistic values of 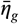, specialists can become locally quite rare, but maintain a global majority. This raises an important question: could we estimate global symbiont diversity reasonably well based on specialists alone?

#### Smaller samples have more false specialists

Real-world ecological datasets often contain a high proportion of apparent specialists, but also have a high proportion of missing hosts and missing links, and so usually under-estimate symbiont host range (*52*). Therefore, many of these “specialists” are, in fact, generalists misrepresented by an incomplete sample of both hosts and host associations. In some cases, the proportion of false specialists can be quite high, especially if networks are sampled incompletely (**Figure 4**).

**Figure 4.**
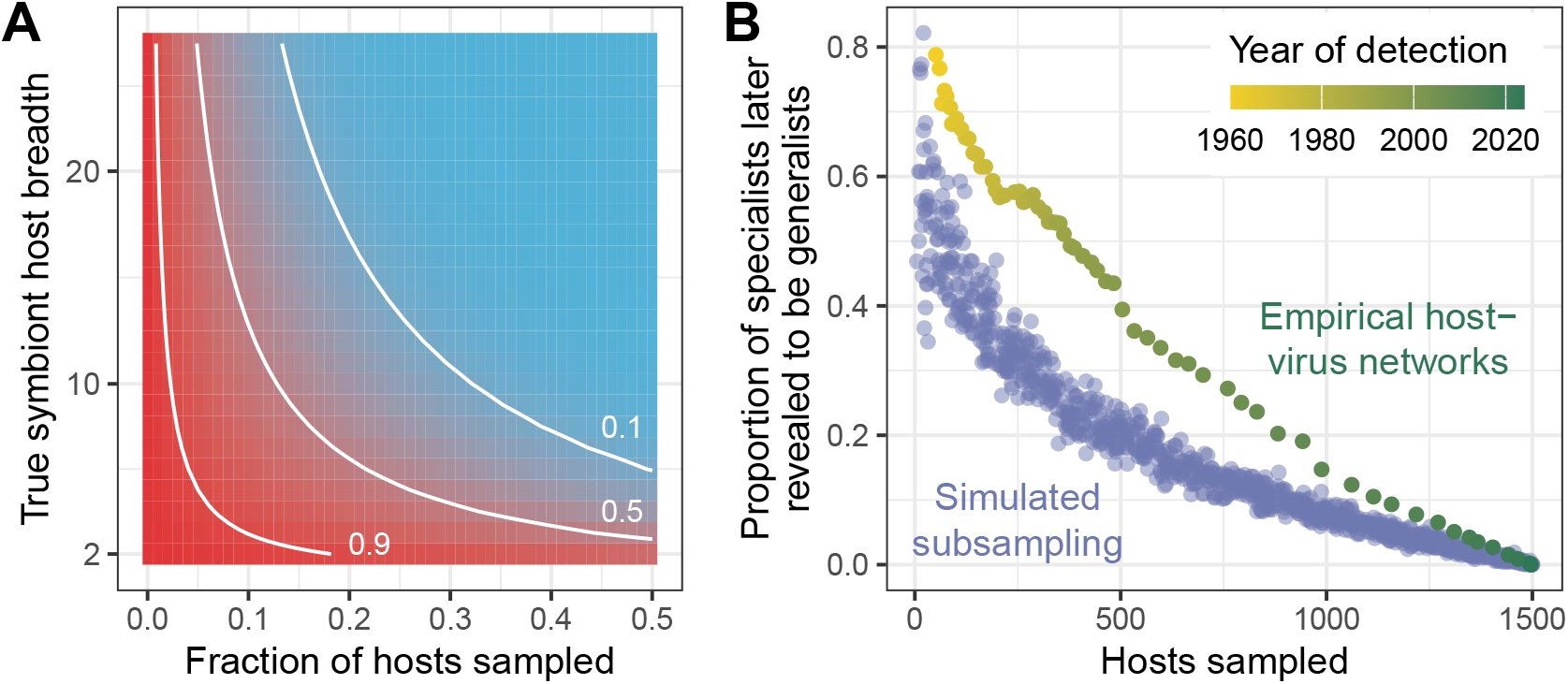
False specialists are common even in well-sampled networks. (A) The expected probability of a given symbiont being represented as a false specialist (red: higher values; blue: lower values; isoclines give reference points), given in equation 13 as a function of the proportion of hosts sampled and the true host range. (B) Scaling between hosts sampled and the the proportion of observed specialists that are false specialists, based on the currently-known global network of mammalian viruses. Simulations (lilac) are based on 1,000 random draws of hosts and all associated viruses; empirical networks (yellow to green) are based on the subset of interactions detected by a given year, based on the first known publication, collection, or data release date (*54*).

Even in a complete sub-sample of *h* hosts, all associated symbionts, and no missing links, a symbiont with *k* true hosts has a probability of having one known host given by equation 1:

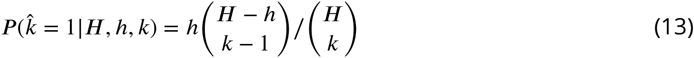

Meanwhile, the probability that the same symbiont is missing from the sample entirely is given by:

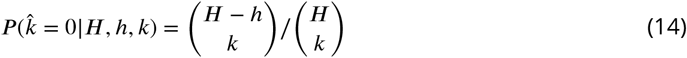

Therefore, in small subgraphs, most symbionts will not be counted — but of those that are, a very high proportion of generalists will be falsely recorded as specialists. For example, consider a tapeworm with *k* = 10 mammal hosts out of a total of *H* ≈ 6, 000 hosts: in a subsample of 10% of hosts (*h* = 600), there is a ∼34.8% chance the parasite will not be detected, but if it is, there is a 59.5% chance it will be recorded as a false specialist. In a sample of just 1% of hosts (*h* = 60), there is a 90.4% chance the parasite will not be detected, but if it is, the probability that 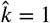 rises to 95.6%. Therefore, for symbioses where most hosts are still unsampled and most symbionts are believed to be undiscovered — for example, all microbial pathogens and most macroparasites — it is safe to assume that the host ranges of many symbiont species are substantially under-estimated. In even smaller samples, such as some of the site-specific datasets widely used in ecology, it would be reasonable to conclude that most observed specialists are false specialists.

Therefore, even when the majority of symbionts are specialists, extrapolating a lower bound on global diversity based on observed “specialist” symbionts may lead to dramatic over-estimation, compared to a balanced approach that accounts for the full distribution of host range.

### Are most endangered species symbionts?

Previous work has suggested that coextinction might be a substantial contributor to present-day biodiversity loss, or could even account for a “silent majority” of total biological extinctions (*5, 6, 62*). On one hand, symbionts frequently outnumber their hosts in terms of total species richness, especially in the case of hyperdiverse groups like beetles or microbes. On the other, symbiont coextinction *rates* are by definition equal to or *lower* than host extinction rates, except in cases where symbionts are disproportionately (non-randomly) associated with high-vulnerability hosts.

Based on the scaling relationships that we have described here, we propose three hypotheses about symbiont coextinction. First, we suspect that coextinction *rates* are frequently overestimated, especially by studies that simulate coextinction based on interaction datasets collected at the level of a single ecological community. Smaller samples are not only full of false specialists, but more generally prone to under-estimating symbiont host range (**Figure 5A**). However, there is also a hard constraint imposed by the linear scaling of specialist coextinctions, and the estimated host extinction rate is probably a greater source of uncertainty (**Figure 5B**).

**Figure 5.**
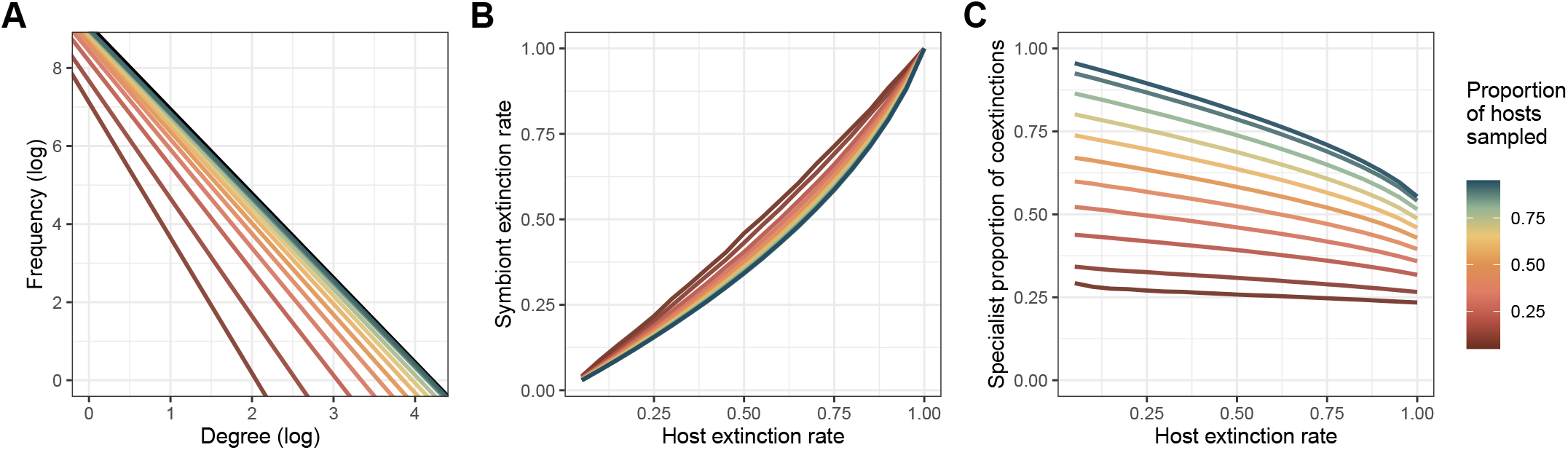
Specialists drive global coextinction risk. To simulate real-world sampling, subgraphs were drawn from the global vertebrate parasite network (see **Methods**) at levels of host sampling ranging from 5% to 95%, with 100 simulations at each level for degree distributions and 500 simulations for coextinction curves, and 20 simulated host extinction rates ranging from 5% to 100%.

Second, despite over-estimated coextinction rates, the total *number* of endangered specialists is likely to be high. Some symbioses are simply not globally hyperdiverse (for example, clownfish and anemones); but for those that are, it is unlikely that any given host group will somehow be immune to the ongoing global loss of biodiversity. For any symbiosis in which the average host has at least one specialist symbiont (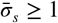), endangered symbionts will outnumber endangered hosts by default. For example, Larsen and colleagues estimate that every species of arthropod hosts “at least one species-specific mite, nematode, apicomplexan protist, and microsporidian fungal species, and a mean of 10.7 bacterial species” (*42*). In this scenario, arthropods’ parasites and pathogens could easily be one of the most diverse groups on Earth, and by numbers alone, they would account for the majority of endangered species.

Finally, in most hyperdiverse symbiont groups, specialists are likely to account for the global *majority* of coextinctions. Generalists are always less vulnerable to coextinction, but especially at lower host extinction rates (**Figure 5C**). At planetary scales, we should expect that specialists not only face the highest coextinction risk, but also account for the majority of endangered species. Today, roughly 1.6% of free-living species face extinction risk from climate change (*63*), and 1.4% face extinction risk from agriculture-related habitat loss (*64*). With roughly 3°C of warming projected by the end of the century, extinction risk from climate change could reach 5-6% globally. At these “low” levels, the number of potential specialist coextinctions across different forms of symbiosis could be staggering. For many groups — especially microbes and invertebrates — most of these specialists are unlikely to be discovered and described before they go extinct (*31*).

## Conclusion

One of the earliest controversies in the study of ecological networks was a debate between Cohen & Briand (*65*), who argued that the number of links in food webs scaled linearly with the number of species; and Martinez (*66*), who held that connectance was constant across networks. The remaining discrepancies between these predictions, and novel data, led Brose and colleagues (*67*) to suggest that the number of such interactions scaled with the number of species following a power law. This relationship was then called into question by MacDonald and colleagues (*68*), who refined the scaling relationship to account for structural constraints coming from the number of species. Each of these studies took an incremental step towards better ecological theory, but not for theory’s sake alone. We can only conserve what we know exists: knowing how species interactions scale with species richness helps us measure and monitor an important facet of biodiversity.

More than forty years after ecologists began to debate the existence of these scaling laws, it would be natural to assume that the question “How do ecological networks scale?” would be mostly answered. However, our results suggest that the macroecology of symbiosis is still in its infancy. The study of small, uncharismatic, and “mostly harmless” organisms has always received less attention, and fewer resources, than research focused on charismatic, free-living, easily-observed species with obvious value to, and interactions with, humans (*8, 69, 70*). Nevertheless, as we show here, symbionts — whether beneficial, neutral, or harmful to their host — are likely to account for the majority of global biodiversity, and may well suffer some of the greatest losses from human alterations of the biosphere. The widespread loss of these species, and the millions of ecological interactions they allow to exist, will have consequences for ecosystem health and resilience that are all but impossible to predict — but knowing what exists today is a good first step. The task of describing and protecting symbiotic species and their interactions is a global one, and will continue for decades to come; so too will the need to pursue theory relentlessly and on all fronts. As with undetected symbionts, we do not know how many laws of macroecology are left to discover (or disprove), and how they could help us protect planetary biodiversity.

## Materials and Methods

### Data sources

No original data were generated in this study.

#### Global data

Vertebrate-virus association data (shown in Figures 3, 4, S1, and S6 and Table S2) were taken from the VIRION dataset and accessed using the virionData R package (*54*). We subset these data to interactions that are established using high-confidence detection methods (PCR and viral isolation), and exclude humans as a host due to their oversampling.

Vertebrate-helminth association data (shown in Figures 5 and S5) were taken from the Natural History Museum’s Host-Parasite Database and originally accessed using the helminthR R package (*57*). Here, we reuse a version of this dataset we previously subjected to basic taxonomic cleaning (*31*). In Figure 5, we subset this dataset to vertebrate parasites; and in Figure S3, we subset this dataset to nematode parasites of mammals.

#### Community-level datasets

A compendium of 318 community-level species interaction datasets (used in Figures 2, S2, and S4 and Table S1) were downloaded from the Web of Life database (*71*). We exclude food webs and limit our analysis to pairwise interactions, which include plant-floral visitor (n = 174), host-parasite (n = 51), plant-seed disperser (n = 34), anemone-fish (n = 17), plant-ant (n = 5), and plant-herbivore (n = 4). Throughout, we limit our analyses to networks with at least five hosts and five symbionts (n = 253 of 281 total datasets).

#### Supplementary sources

In the supplement, we adapt Figure 1 from (*22*), which reused three interaction datasets from the literature: plant-seed disperser associations in Kenya (*72*); plant-mycorrhizal fungi associations in Japan (*73*); and plant-floral visitor associations in Illinois, USA (*74*). For the final panel, we use the updated dataset of mammal-nematode associations described above (*31, 57*).

### Analyses

All analyses were conducted in the `R` statistical language. Statistical analyses were conducted using the base stats package. Network metrics were calculated using the `bipartite, codependent`, and `igraph` packages (*22, 75, 76*). In the section “How many symbionts are there?” we use the `iNEXT` package to estimate the diversity of viruses based on the observed network (*49*). Visualizations were created using the `ggplot2` package (*77*).

## Data and Code Availability

All data and code are available on Github (www.github.com/carlsonlab/straightlinewasalie).

## Acknowledgments

This project has benefited from feedback and discussion with several colleagues over almost a decade, including Greg Albery, Laura Alexander, Allison Barner, Tad Dallas, Andrew MacDonald, Phillip Staniczenko, Zachary Susswein, and most of all Shweta Bansal. We used color schemes based on the art of Lily Paris West, and the LaPreprint template crated by Mikkel Roald-Arbøl, and thank both for helping make our manuscript more beautiful.

JBY thanks Mr. Stephen Sondheim, for the reminder that one-sided graphs are a poor representation of the most interesting species interactions: it takes two. CJC also thanks Mr. Sondheim, for the reminder that you have to finish the hat; and Dr. Stuart Sidney, for making calculus even more fun than Tom Lehrer made arithmetic. CJC finally dedicates this study to Dr. Carole Wade, without whom the work would never have been started, let alone completed.

## Author contributions

All authors contributed to the conceptualization of the study, the analysis of the data, and the writing and editing of the manuscript.

## Supporting Information

**Table S1.** ANOVA for data shown in Figure 2C.

|  | Df | Sum Sq | Mean Sq | F value | *Pr(> F)* |
| --- | --- | --- | --- | --- | --- |
| Connectance |  |  |  |  |  |
| Interaction type | 2 | 1.57 | 0.78 | 44.51 | $< 10^{-6}$ |
| Full network or generalists | 1 | 1.07 | 1.07 | 60.53 | $< 10^{-6}$ |
| Type × network | 2 | 0.03 | 0.02 | 0.90 | 0.41 |
| Residuals | 486 | 8.56 | 0.02 |  |  |
| Modularity |  |  |  |  |  |
| Interaction type | 2 | 2.01 | 1.01 | 59.34 | $< 10^{-6}$ |
| Full network or generalists | 1 | 0.69 | 0.69 | 40.54 | $< 10^{-6}$ |
| Type × network | 2 | 0.08 | 0.04 | 2.26 | 0.11 |
| Residuals | 486 | 8.23 | 0.02 |  |  |
| Nestedness |  |  |  |  |  |
| Interaction type | 2 | 3.08 | 1.54 | 61.34 | $< 10^{-6}$ |
| Full network or generalists | 1 | 0.59 | 0.59 | 23.42 | $< 10^{-5}$ |
| Type × network | 2 | 0.05 | 0.03 | 1.07 | 0.34 |
| Residuals | 486 | 12.20 | 0.03 |  |  |

**Table S2.** Estimates of viral diversity per host (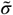) based on metagenomic samples (*40, 41*), and average viral host range (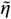) for all viruses in the VIRION database (*54*) and just those that are ICTV-ratified.

| Virus family | $\tilde{\sigma}$ | $\tilde{\eta}$ (all) | $\tilde{\eta}$ (ICTV) | $\tilde{\sigma}/\tilde{\eta}$ (all) | $\tilde{\sigma}/\tilde{\eta}$ (ICTV) |
| --- | --- | --- | --- | --- | --- |
| DNA viruses |  |  |  |  |  |
| Adenoviridae | 8.5 | 1.8 | 3.5 | 4.8 | 2.4 |
| Orthoherpesviridae | 5.5 | 1.7 | 3.9 | 3.3 | 1.4 |
| Paramyxoviridae | 5.5 | 3.2 | 4.2 | 1.7 | 1.3 |
| Parvoviridae | 7 | 1.8 | 2.3 | 3.8 | 3 |
| Polyomaviridae | 2.5 | 1.2 | 1.1 | 2.1 | 2.3 |
| RNA viruses |  |  |  |  |  |
| Astroviridae | 13 | 2.6 | 3 | 5 | 4.3 |
| Coronaviridae | 3 | 4.7 | 8 | 0.6 | 0.4 |
| Picobirnaviridae | 60 | 1.5 | 8 | 41.3 | 7.5 |
| Picornaviridae | 2.5 | 2.5 | 4.5 | 1 | 0.6 |
| Rhabdoviridae | 1 | 3.5 | 4.9 | 0.3 | 0.2 |
| Sedoreoviridae | 1.5 | 3.7 | 7.1 | 0.4 | 0.2 |

**Figure S1.**
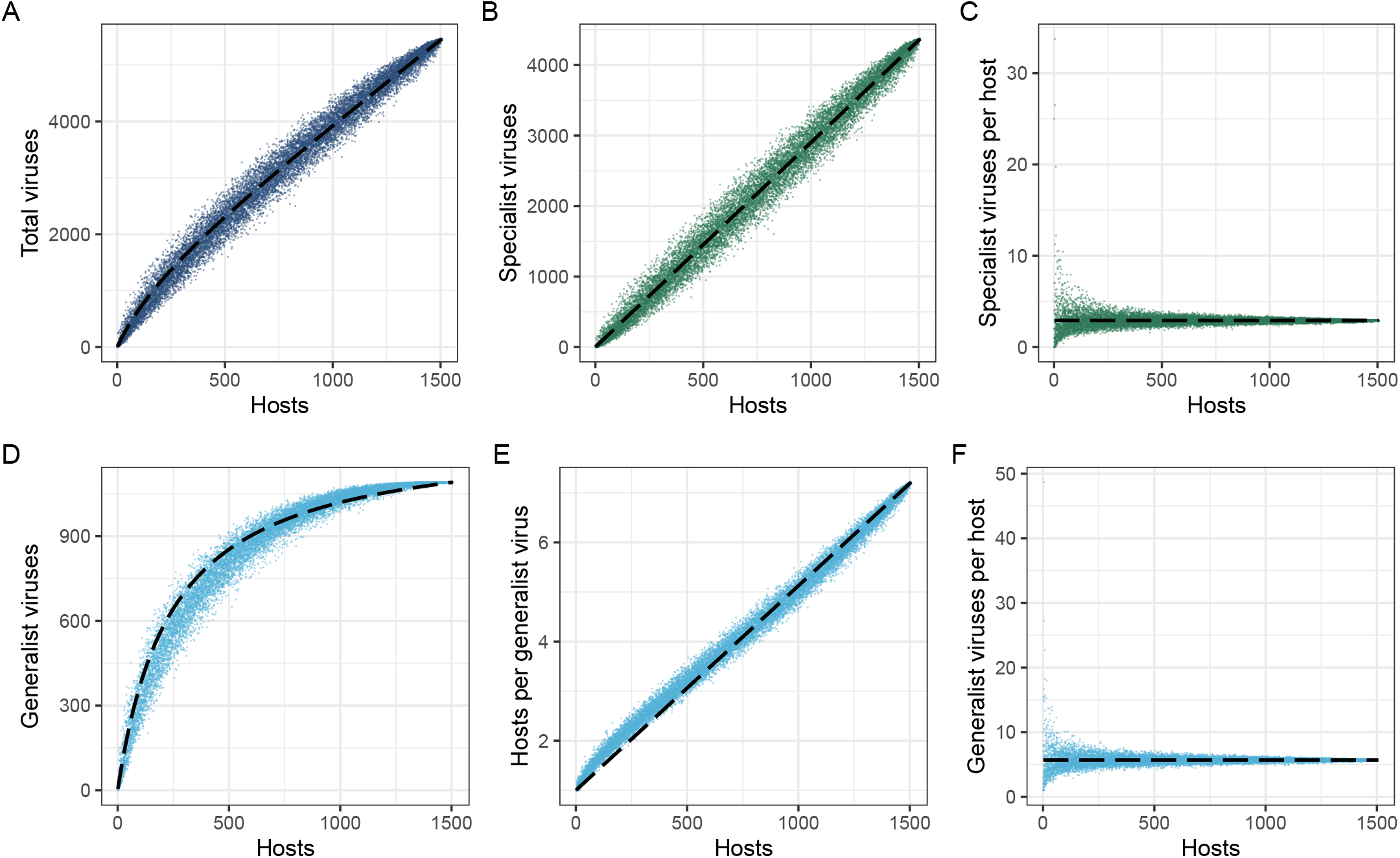
Sub-linear scaling can be approximated based on three network-level parameters. Dotted lines show the approximation estimated in equation 9, split into its component parts. Points show 1,000 samples of the mammal-virus network with a randomly selected number of hosts (*54*).

**Figure S2.**
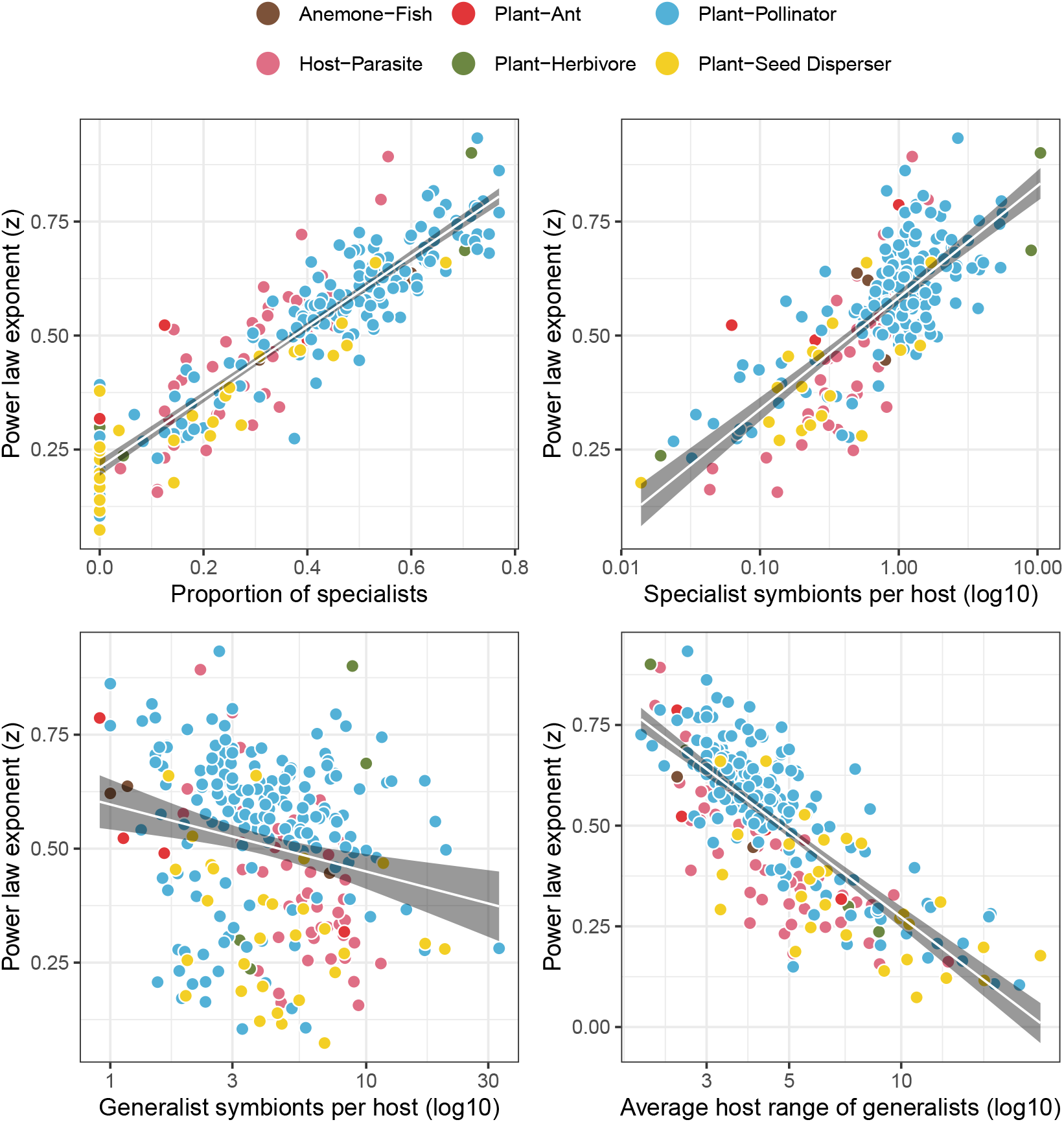
The specialist-generalist split explains “power law” scaling. Estimated power law exponents (z) compared to the parameters that explain equation 9, compared across 253 real ecological networks (see **Methods**).

**Figure S3.**
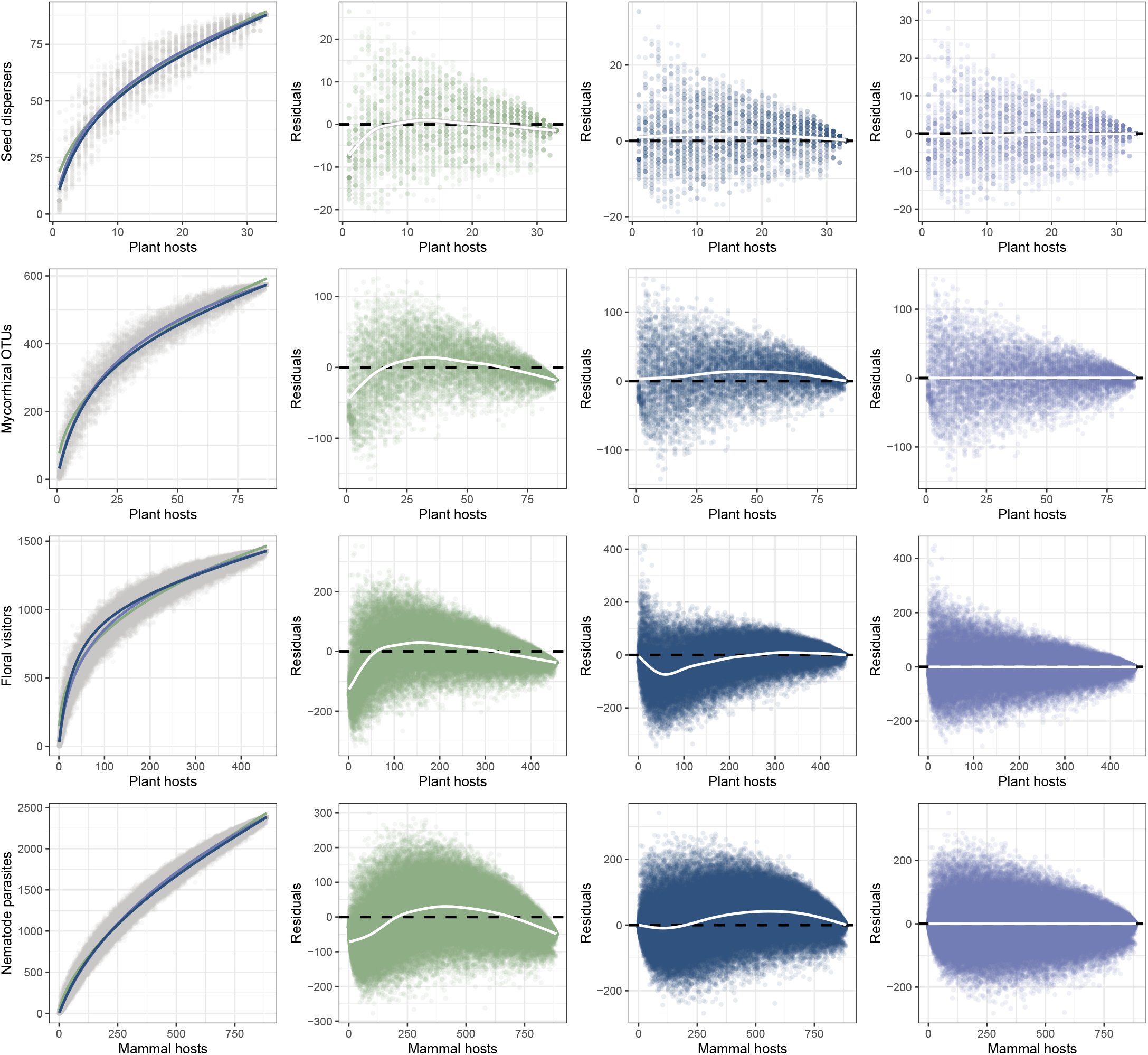
First-principles solutions compared to power law estimates. Datasets are reanalyzed from Figure 1 of (*22*), with estimates based on a power law (green), our three-parameter approximation (blue), and the Koh-Colwell method (purple).

**Figure S4.**
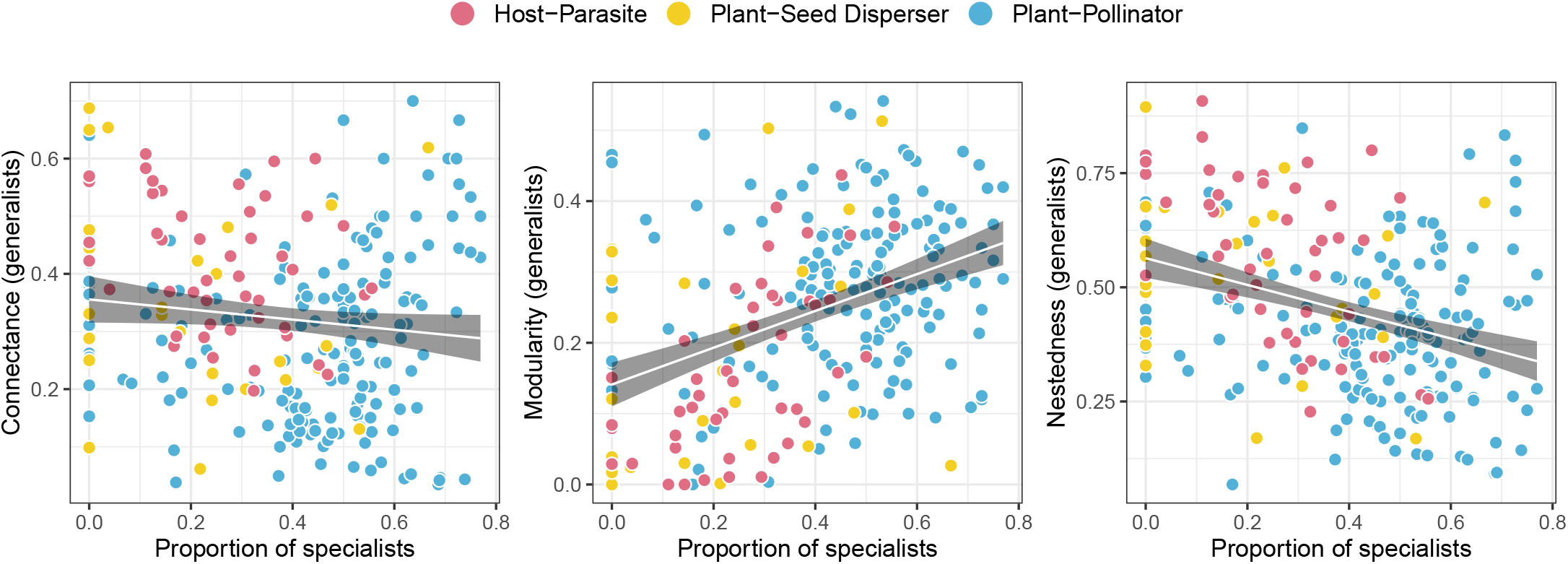
Specialists are weakly correlated with host-generalist network architecture. See caption to Figure 2.

**Figure S5.**
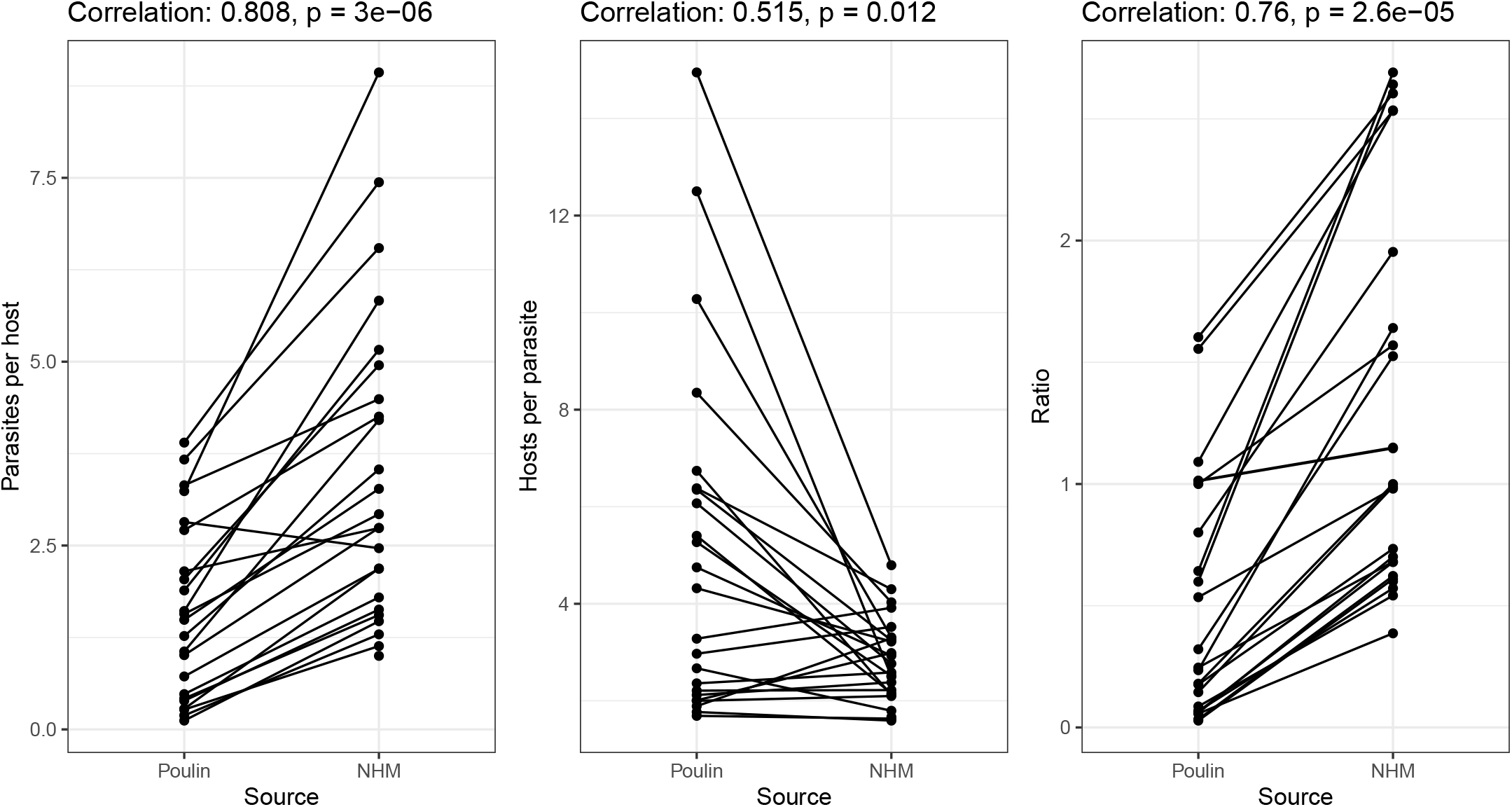
Global parameters of the host-parasite network have changed over time. Each point is an estimate for a combination of vertebrate host class and helminth taxonomic group, e.g., nematode parasites of birds. Data are adapted from (*19*) and (*31*).

**Figure S6.**
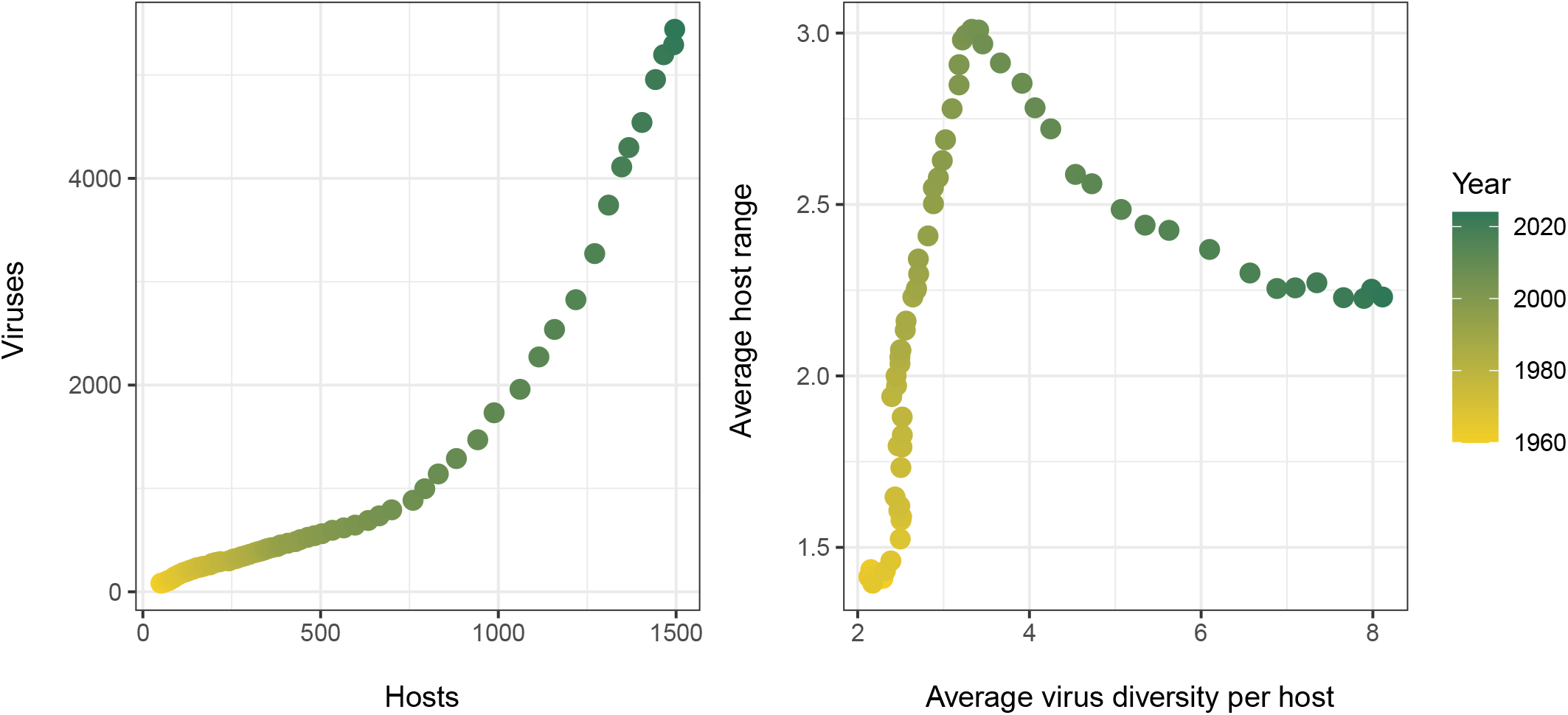
The observed host-virus network through time. Data reflect the earliest known year of association (*54*).

